# Simulation-based spatially explicit close-kin mark-recapture

**DOI:** 10.1101/2025.05.31.656892

**Authors:** Gilia Patterson, Claire K. Goodfellow, Nelson Ting, Andrew D. Kern, Peter L. Ralph

## Abstract

Estimating the size of wild populations is a critical priority for ecologists and conservation biologists, but tools to do so are often labor intensive and expensive. A promising set of newer approaches are based on genetic data, which can be cheaper to obtain and less invasive than information from more direct observation. One of these approaches is close-kin mark-recapture (CKMR), a type of method that uses genetic data to identify kin pairs and estimates population size from these pairs. Although CKMR methods are promising, a major limitation to using them more broadly is a lack of CKMR models that can deal with spatial heterogeneity both in population density and sample effort. We introduce a simulation-based approach to CKMR that uses spatially explicit individual-based simulation in concert with a deep convolutional neural network to estimate population sizes. Using extensive simulation, we show that our method, CKMRnn, is highly accurate, even in the face of spatial heterogeneity, and demonstrate that it can account for potential confounders such as unknown population histories. Finally, to demonstrate the accuracy of our method in an empirical system, we apply CKMRnn to data from a Ugandan elephant population, and show that point estimates from our method recapitulate those from traditional estimators but that the confidence interval on our estimator is reduced by approximately 30%.

## Introduction

Population size is one of the most important pieces of information that informs the conservation and management of wildlife populations, but it is often challenging and expensive to estimate. One of the most promising sources of information about population size is genetic data. Genetic material can be relatively easily collected from sources such as hair, scat, or hunter harvests, and it is increasingly affordable to generate genotype information from this material. Thus developing accurate statistical methods to use genetic data has the potential to improve monitoring and conservation efforts for many species.

One of the most commonly used methods for estimating population size from genetic data is genetic capture-recapture, which uses genotype information sampled non-invasively, for example, from hair or scat, to identify when the same individual has been captured multiple times and then estimates census population size from these recaptures [27, 29]. Genetic capture-recapture models can account for many of the complexities of real populations, such as unequal capture probabilities due to behavioral responses to trapping or differences in the amount of DNA shed by individuals [27]. More recently, these models have been extended to use spatial information. Spatially-explicit capture-recapture models can account for the spatial distribution of sampling locations on the landscape, ecological variables that impact the movement of individuals, and variation in population density across the landscape [13, 14]. Genetic capture-recapture and spatially explicit genetic capture-recapture have been used to effectively monitor many species including grizzly bears from hair snares [4], red deer from scat [12], and fishers and grey foxes from hair traps [19].

Close-kin mark-recapture (CKMR) is a promising, recently developed type of method that uses genetic data to identify pairs of close kin among a set of sampled individuals, and then uses these pairs in a way conceptually similar to recaptures to estimate abundance [6, 23]. Because CKMR does not require recaptures, it has the potential to improve monitoring for many of the species for which genetic capture-recapture models cannot be used or do not provide good estimates. These include systems that require lethal sampling and systems with elusive, hard to capture individuals. CKMR models have been used to estimate abundance in several species, including southern bluefin tuna [5], white sharks [21], thornback rays [40], brook trout [40], speartooth sharks [30], flying foxes [26], caribou [28], bearded seals [39], and lemon sharks [38]. However, a major limitation to using CKMR methods more broadly is that spatially explicit methods that account for population structure and spatial biases in sampling are limited, with only one recently published method available [34]. When dispersal is limited (i.e., when spatial population structure exists), kin pairs are expected to be found more closely together, and so when the landscape is sampled more intensely in some areas than others, the number of sampled kin pairs is larger than expected. This results in downwardly-biased estimates of population size when non-spatial methods are applied [9].

Current CKMR methods (with the exception of Conn [8]) are generally based on an analytic likelihood of observing a given kin relationship for each pair of individuals in the sample. The basic likelihood models for parent-offspring and half-sibling pairs are described in Bravington, Skaug, and Anderson [6]. Most applications of CKMR incorporate more complicated population dynamics into these likelihood models by necessity. For example, Lloyd-Jones et al. [26] added a parameter for recent population size trend to estimate both the trajectory of the population and abundance in flying foxes. Swenson et al. [38] developed models that incorporated uncertainty in aging, fluctuations in population size, and intermittent breeding. Finally, Sévêque et al. [34] accounted for spatial population structure when relative density across the landscape is known and dispersal is independent between individuals.

Adding spatial information to likelihood methods becomes much more difficult when models of movement and dispersal are complex, both because formulating an appropriate likelihood becomes challenging and because computing the likelihood becomes computationally intensive. Simulation-based inference provides a promising way of overcoming these challenges. Simulation-based methods do not require a likelihood and the complexity of the model is limited only by the ability to simulate reasonable approximations to the true population dynamics [10]. Recently, simulation-based inference models have begun to be applied to non-spatial close-kin mark-recapture and show great promise. Conn [8] implemented individual-based simulations and used observed counts of kin pairs grouped by age classes with Approximate Bayesian Computation (ABC) to infer posterior distributions of abundance and survival. This approach accounted for non-independence of pairs of individuals and dealt with the inability to distinguish half-sibling relationships from aunt-niece or grandparent-grandchild, two difficulties with previous likelihood-based methods.

In this paper, we develop a novel simulation-based spatially explicit close-kin mark-recapture method, named CKMRnn. We create synthetic “images” of kin pairs and sampling intensity across the landscape that compactly encode the spatial information available for the system, then train a deep convolutional neural network (CNN) on simulated images to estimate population size. Similar simulation-based methods utilizing CNNs have been demonstrated to be effective at inferring population genetic parameters from genetic data [15], and more specifically, for spatial population genetic parameters such as density and dispersal [35, 36, 37].

We first test CKMRnn on simulated populations with limited dispersal, spatially-biased sampling, and varying trends in population size and demonstrate that our method is both accurate and robust. We then apply CKMRnn to estimate the population size of African elephants (*Loxodonta africana* and *L. cyclotis*) in Kibale National Park in Uganda using data from dung samples collected by Goodfellow et al. [17]. Like many populations of conservation concern, the Kibale elephant population is challenging to monitor and has strong spatial patterns in kin pairs across the landscape that could lead to bias when using traditional, non-spatial CKMR methods. This makes Kibale elephants a good system to test the feasibility of our method. We find that CKMRnn agrees with traditional capture-recapture point estimates of elephant population size, but provides a 32% smaller confidence interval.

## Methods

### CKMRnn workflow

The first step of the CKMRnn workflow is to process the empirical data to create a collection of images summarizing the observed kin pairs and sampling effort. The images are all of the same dimension and show the geographic region of interest. One image summarizes the intensity of the sampling effort across the region as a heatmap and the other images show the observed kin pairs and/or recaptures by connecting their sampling locations with line segments (see Figure 1). The precise details and number of these images may differ depending on the system. For instance, “sampling effort” could be measured as the amount of time spent searching in each section of a study landscape, as the number of samples found in each section, or as a variety of other metrics. The total number of images will depend on how many types of close kin and/or recaptures are used. For instance, one could apply the method to *only* mark–recapture data, with only one image connecting the mark/recapture locations; while another situation might have separate images for full siblings, half siblings, parent-offspring pairs, etcetera. In our application to elephants, we made these images by first projecting the GPS coordinates of each sample onto a rectangular surface using GIS tools in R, then creating images using the Python Image Library.

**Figure 1:**
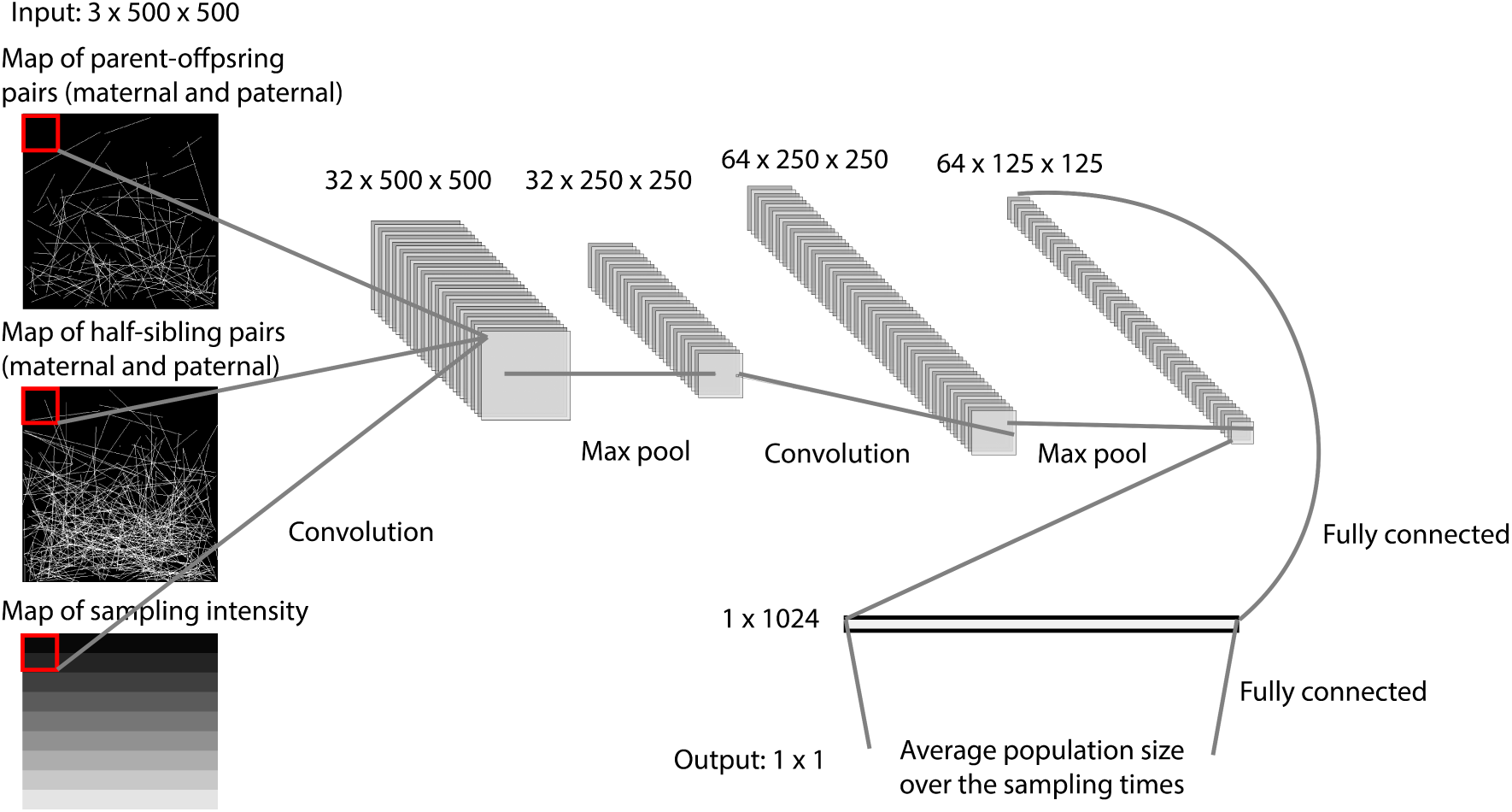
CKMRnn’s convolutional neural network architecture.

The second step is to develop a spatially explicit individual-based simulation of the system in question, including the empirical sampling scheme. In our examples we developed simulations using the population genetic simulation software SLiM [20], building on the spatial simulation methods described in [7]. The dynamics of the simulation are based on previous knowledge of the population of interest. The simulation does not need to perfectly match the empirical population, and we can account for uncertainty in some of the parameters of the simulation by choosing a reasonable range of values, similar to a prior distribution, and then simulating across this range.

The third step is to generate training data and train the neural network. Training data is produced by running many simulations with a range of population sizes (and possibly other parameter values), then processing each simulated sample to produce images of the same dimensions and summarizing the same information as the images created for the empirical data. The convolutional neural network (CNN) is then trained on this training data to estimate population size from images.

The final step is to estimate population size and uncertainty for the empirical population. The point estimate of population size is obtained by passing the images derived from the empirical data to the trained network. Then, a distribution of parametric bootstrap replicates is generated by running many simulations with population size set to the point estimate and passing the resulting images for each simulation through the trained network. A confidence interval is then computed from this distribution of estimates.

We describe these steps in more detail in our examples below. The code for the CKMRnn workflow is available in the CKMRnn GitHub repository (https://github.com/giliapatterson/CKMRnn).

### Simulation tests

We start by evaluating the performance of CKMRnn using simulations. Our simulations closely follow those used by Conn et al. [9], who used simulations modeled on bearded seals to demonstrate that non-spatial CKMR can be biased in the presence of spatial structure. We implemented the simulation using the simulation software SLiM [20], incorporating age-dependent survival and reproduction, limited dispersal, and lethal sampling. The simulation is stochastic and individual based and takes place in continuous space.

The simulation proceeds on a homogeneous 10 × 10 square landscape (the units are arbitrary, but taking density estimates from Fuirst et al. [16], one unit might roughly correspond to 10km). The population is initialized with *N*_0_ individuals, with an even sex ratio, where initial individual locations are chosen uniformly across the landscape. *N*_0_ is varied between simulations to obtain different population densities. The initial ages for individuals are assigned from the stable age distribution based on a non-spatial Leslie matrix model with the same survival and reproduction parameters as our simulation. Each timestep of the simulation represents one year, and proceeds through reproduction, survival, and dispersal. We simulated 40 years of burn-in, then recorded and sampled from all individuals that died in years 40 through 60.

In each time step, first, fertile females mate and reproduce. A female of age *a* is fertile with probability 1*/*(1 + exp(−1.264(*a* − 5.424))). If a female is fertile, she chooses a mate from all males within a radius of 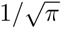. Older males are more likely to be chosen: the probability that a male of age *a_m_* within the circle is chosen as a mate is proportional to 1*/*(1 +exp(−1.868(*a_m_* − 6.5)))). Numerical constants in these expressions were taken from Conn et al. [9]. If at least one male is within the circle, the female produces one offspring, and the offspring’s location is set to that of the female’s.

Next, each individual of age *a* dies with probability

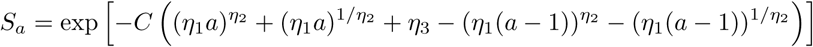

where *C* = 1.111, *η*_1_ = exp(−2.904), *η*_2_ = 1 +exp(0.586), and *η*_3_ = exp(−2.579). The parameters *η*_1_, *η*_2_, and *η*_3_ were again taken from Conn et al. [9]. The parameter *C* was calculated using the Leslie matrix model and chosen so that the population size stays constant. We also introduced a linear population trend into the simulation by multiplying survival probabilities by 1.01 or 0.99.

Finally, each individual moves to a new location chosen from a bivariate Normal distribution centered on the original location with standard deviation *σ* = 1. This means that all individuals move on average a distance of 1 unit, or roughly 10 km, each year.

To simulate sampling, we divided the landscape into a grid of one hundred 1 × 1 squares, assigned a sampling intensity *I_i,j_* to each grid cell, and then took a random sample of 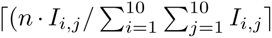 individuals from the *i, j*^th^ grid cell, for each cell, where *n* is the total sample size. In all sampling plans, sampling intensity was constant across rows and increased linearly in columns, so that *I_i,j_* = 1 + *α*(*i* − 1)*/*10 for some *α*. We implemented three sampling plans: (1) uniform, where all intensities were equal (*α* = 0); (2) medium spatial bias, where the bottom row of the landscape was 15 times more likely to be sampled than the top row (*α* = 15); and (3) high spatial bias, where the bottom row of the landscape was 30 times more likely to be sampled (*α* = 30).

We generated two sets of simulated data, one with constant population size and one with varying population trends. For the constant size dataset, we ran 2601 simulations with initial population sizes between 5000 and 20000. For the varying trends dataset, we ran 2601 simulations with each of the three trends (constant, increasing, and decreasing). We recorded final and average population sizes for each simulation. For each simulation, we sampled *n* = 2000 individuals using uniform, medium-bias, and high-bias sampling plans. The constant size dataset had 7803 total simulated samples and the varying trends dataset had 23409 total samples. For each sample, we created three 500 × 500 pixel images. The first is a map showing relative sampling intensity across the landscape (the 10 × 10 image of *I_i,j_*). The second two show half-sibling and parent-offspring pairs, respectively, with pairs connected by line segments.

The neural network architecture is summarized in Figure 1. The first layer is a convolutional layer with 3 input channels, one for each 500 × 500 image, and 32 output channels. This layer has a kernel size of 6 and padding of 3. It performs a 2d convolution separately on each input channel (image) and then sums the result. Each of the 32 output channels is a 500 × 500 matrix that is the result of a separate convolution. Each matrix is passed through a ReLU layer that converts negative entries to 0 and then through a max pooling layer with a kernel size of 2. The max pooling layer divides each of the 32 500 × 500 matrices into 250 2 × 2 squares and keeps only the maximum value in each square, resulting in 32 250 × 250 matrices. The next three layers are another set of convolution, ReLU, and max pool layers. This time, the convolution layer has 32 input channels and 64 output channels and so the output of these 3 layers is 64 125 × 125 matrices. These matrices are flattened into a single, 1 × 10^6^ vector and then passed through a fully connected layer with 1024 output nodes. The resulting 1 × 1024 vector is then passed through a ReLU layer and a dropout layer. The dropout layer sets 0.2 of the elements of the vector to 0. The final layer is a fully connected layer that outputs a single number, the estimated population size.

We trained two neural networks, one on the constant size dataset and one on the varying trends dataset. For each network, one half of the simulations were used for training, one quarter for validation, and one quarter for testing. We trained the network to minimize mean-squared error loss using the Adam optimizer and batches of size 64. Training was run for 20 epochs.

### Estimation of population size in African elephants

We next used CKMRnn to estimate population size of African elephants in Kibale National Park in Uganda using the genetic samples collected by Goodfellow et al. [17]. The dataset contains 256 samples collected from dung piles across Kibale (Figure 2A). These samples were collected on 49 days between November 2020 and August 2021 from locations around the park where elephants were known to spend time. The samples were genotyped at 14 microsatellite loci and these genotypes were analyzed with GenAlEx v. 6.51b2 and MLRelate [24] to identify unique individuals and parent-offspring pairs [17, 18].

**Figure 2:**
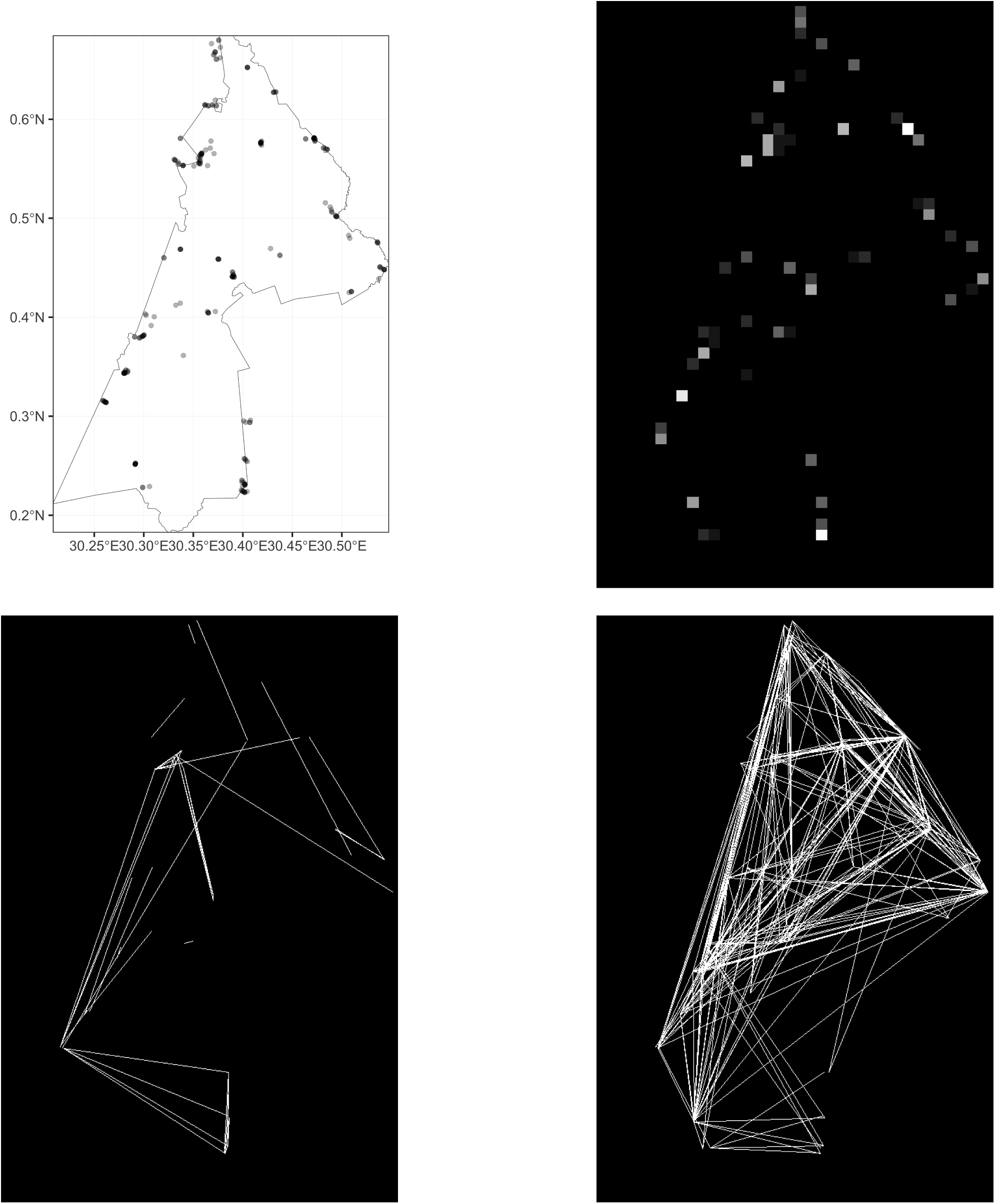
Empirical data for African elephants in Kibale National Park. **(a)** Locations of samples (points) within the park (outline). **(b-d)** Images provided to the neural network, showing **(b)** sampling intensity (pixel lightness is proportional to number of samples in that 1km × 1km pixel); **(c)** recaptures (lines connect original and recapture locations); and **(d)** parent-offspring pairs (lines connect parent and offspring sampling locations).

We first processed the empirical data to create three grayscale images for input to the neural network, one for sample locations, one for parent-offspring pairs, and one for recaptures, as shown in Figure 2. The dimensions of each image were 566 × 837 pixels, and Kibale National park is approximately 38 × 56 km, so this corresponds to 15 pixels per kilometer. To create the images, we pooled all data together ignoring date of collection.

For sampling, we divided the outline of Kibale into a grid of 1 km by 1 km cells, with 771 cells total, and determined the number of samples in each cell. Samples taken from just outside the boundary of the park were assigned to the closest grid cell. The shade of each cell of the sampling image was then proportional to the number of samples, with cells with the maximum number of samples set to white and cells with no samples set to black. Because we do not have information on locations that the field team visited but did not find elephant dung, this image does not represent sampling intensity in the same way as in the simulation tests. It instead encodes information about spatial bias in sampling through the locations of samples.

For recaptures and parent offspring pairs, we first filtered the dataset to retain only the first location for any instances where the same individual was sampled multiple times on the same day. For recaptures, we connected the location of the two captures with a line segment. When an individual was captured more than twice, we sorted the recaptures by time and connected only subsequent pairs: first recapture to second recapture, second to third, and so on.

For parent-offspring pairs, we connected the location of the parent when it was sampled to the location of the offspring when it was sampled. Some of the individuals in parent-offspring pairs were sampled multiple times. For plotting these pairs, we treated each of the samples like a unique individual, and so these pairs are represented by multiple line segments.

To generate training data for the neural network, we developed an individual-based, continuous space SLiM simulation modeled on the elephant population within Kibale National Park. The dynamics and demographic parameters of the simulation model were based on previously published work on forest and savanna elephants [1, 2].

The simulation is set within the boundaries of Kibale National Park and simulated elephants cannot disperse or move beyond the boundary. Each elephant has a location within the park and can move around from year-to-year and day-to-day. Reproduction and survival happen once each year. Mortality is primarily age-based, with elephants living a maximum of 55 years. We varied three parameters of the simulation when generating our training set: population size (*N*), which was drawn from a uniform distribution between 100 and 1500; yearly movement distance (*σ_D_*), drawn from a uniform distribution between 5 km and 10 km; and the radius within which a female chooses a mate (*σ_M_*), also drawn from a uniform distribution between 5 km and 10 km.

The population is initialized with 1.25*N* elephants, half female and half male, with locations chosen uniformly within the boundaries and ages chosen uniformly from 0 to 60. We keep track of the last year a female reproduced, and the “years since last reproduction” for elephants in the initial population were chosen uniformly between 0 and 6.

Each time step of the simulation represents one year, and proceeds through reproduction, survival, and dispersal steps. In the reproduction step, the probability each female is fertile and therefore can reproduce if a mate is available is determined by her age and the number of years since she last reproduced. A female of age *a* and with *y >* 2 years since last reproduction is fertile with probability

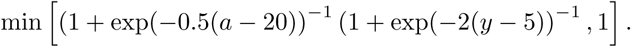

For females with two or fewer years since last reproduction, the probability is 0. For females with at least 6 years since last reproduction, fertility probability starts at 0 and then increases to 1 between ages 10 and 30. For females with 3-5 years since last reproduction, this curve is shifted downward and plateaus at 0.02, 0.12, and 0.5, respectively (Figure 3). Each fertile female chooses a mate uniformly randomly from all males of age greater than 25 that are within a radius *σ_M_* of the female. When a female mates she produces one offspring. To represent the long gestation times of elephants, the offspring is added to a separate population of fetuses, where it remains for two years. If the mother of a fetus is alive after two years, the fetus is transferred to the main population, its age is set to 0, and its location is set to the location of its mother.

**Figure 3:**
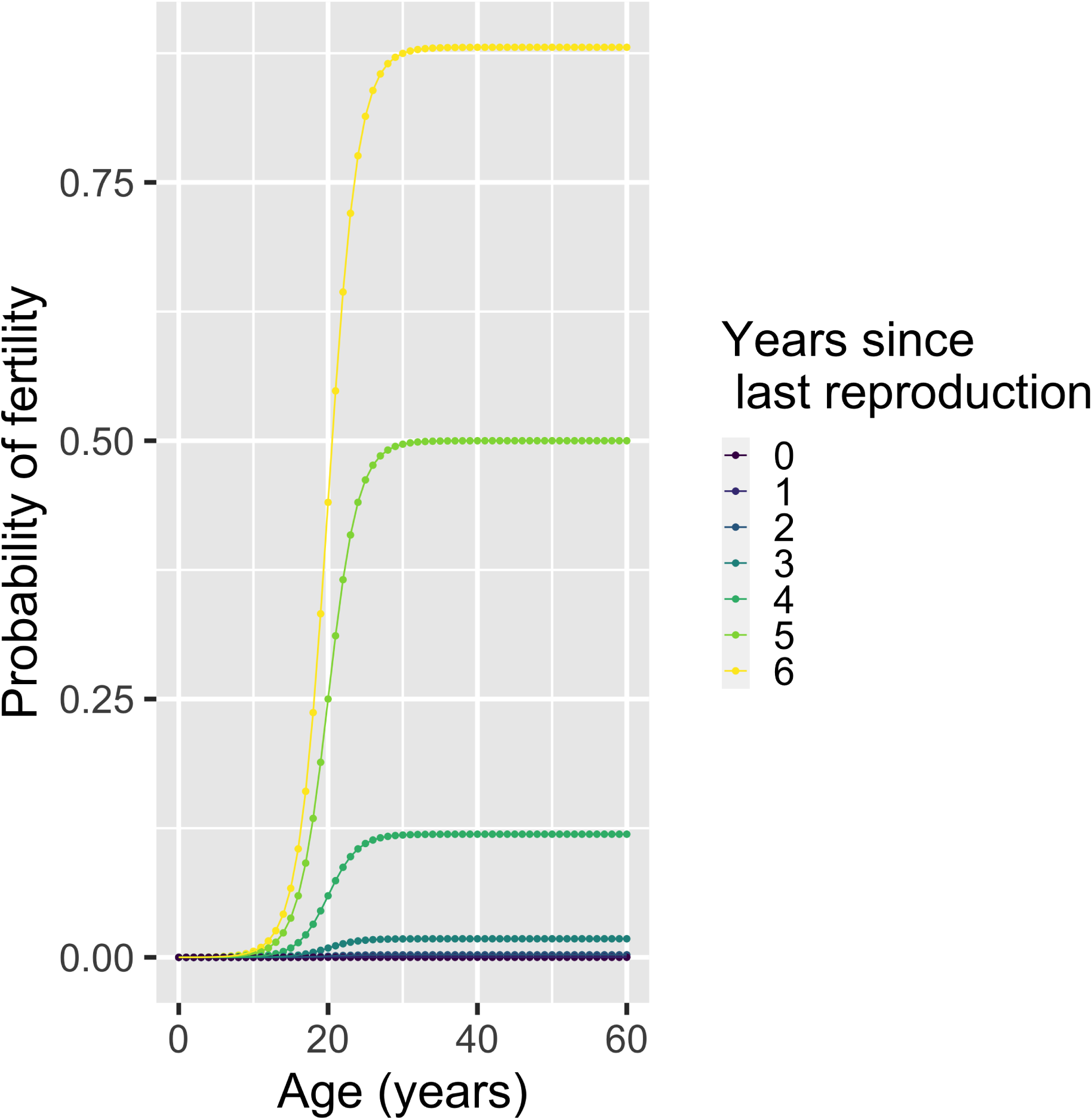
Probability that a simulated female elephant is fertile in a given year, given her age and the number of years since she last reproduced.

Survival is a combination of age-dependent and density-dependent mortality. We took five year probabilities of survival from Armbruster and Lande [1] and converted to one year survival probabilities by taking the fifth root (Table 1). In each time step, we multiplied these fixed survival probabilities for each age by *N* divided by the current population size, ensuring that the population size stays relatively constant around the parameter *N*.

**Table 1:** One year survival probabilities for simulated elephants. Probabilities were calculated by taking the fifth root of five year survival probabilities from Armbruster and Lande [1].

| Age | Female | Male |
| --- | --- | --- |
| 0-4 | 0.871 | 0.871 |
| 5-9 | 0.976 | 0.976 |
| 10-14 | 0.976 | 0.976 |
| 15-19 | 0.979 | 0.979 |
| 20-24 | 0.980 | 0.984 |
| 25-29 | 0.975 | 0.930 |
| 30-34 | 0.975 | 0.924 |
| 35-39 | 0.974 | 0.930 |
| 40-44 | 0.970 | 0.909 |
| 45-49 | 0.910 | 0.652 |
| 50-54 | 0.833 | 0.803 |
| $\geq 55$ | 0.000 | 0.000 |

In the dispersal step, all individuals move around within the bounds of Kibale. New locations for each individual are chosen by sampling a potential new location from a bivariate normal distribution with standard deviation *σ_D_* centered on the current location, checking if the potential new location is within Kibale, and if not, choosing a new potential location. This process is repeated until a location within Kibale is found or until twenty potential locations are drawn, whichever comes first. If no new location is found, the elephant stays in its original location.

Sampling takes place in year 201 after a burn-in period of 200 years. Unlike in other years, when net movement across the year was simulated in a single step, during the dispersal step of the sampling year, 365 days of movement are simulated. Each day, the new locations of individuals are chosen using the same process as for yearly dispersal, but with standard deviation 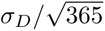. This means that 365 daily movement steps are equivalent to one year of dispersal (except for boundary effects).

To sample our simulated populations, we closely replicated the empirical sampling plan. We first converted the locations and times of empirical samples into SLiM locations and days. For sampling locations, we found the centers of the 1 km by 1 km grid cells that were closest to the locations of the empirical elephant samples, and then used the unique centers as potential sampling locations. For days, we set the first date of sampling to SLiM day 1, and then converted subsequent dates of sampling to SLiM days by calculating the number of days since the first date of sampling. This process led to 53 sampling locations and 49 sampling days. We set the sample size to *n* = 256, matching the empirical sample size.

For each sampling day, a potential sampling location is chosen from the list of empirical locations at random, all individuals within a radius of 1 km from the location are sampled, and then a new potential sampling location is chosen. This process is repeated until either the total number of individuals sampled in the simulation reaches *n* or the number of individuals sampled that day reaches *n/*49. We recorded recaptures, parent-offspring pairs, and locations for sampled simulated individuals.

We ran 10,000 replicate simulations with population sizes drawn uniformly randomly between 100 and 1500 and values of *σ_D_*and *σ_M_*drawn independently from uniform distributions on [5 km, 10 km]. For each simulated sample, we plotted grayscale images of sampling intensity, recaptures, and parent offspring pairs using the same procedure as for the empirical data.

The elephant neural network has a nearly identical architecture to the network in the simulation tests described above. The only difference is that the input size of each of the three images is 566 × 837 instead of 500 × 500 and so the input for the first fully connected layer is a 1 × 1908480 vector instead of a 1 × 10^6^ vector. We trained the elephant network using 7500 simulations and reserved the remaining 1500 for testing. We trained the network to minimize mean-squared error loss using the Adam optimizer and batches of size 10. Training was run for 20 epochs. We implemented the network in the Pytorch package.

## Results

### Simulation tests

To evaluate the performance of CKMRnn on simulated populations, we used the two sets of bearded seal simulations (constant trend and varying trend) to create three tests. First, we measured the performance of CKMRnn in situations where the dynamics of the true population exactly match the training data by training CKMRnn on half of the constant-size set and testing on part of the other half. Second, we measured performance when population trend is misspecified (i.e., there is a trend in the real system not reflected in the simulations) by training on the constant-size set and testing on the varying trend set. Finally, we measured performance when population trend is unknown but not misspecified by training on half of the varying trend set and testing on part of the other half. The last test reflects a general strategy for dealing with unknown aspects of the empirical system: simulate across a “prior” range of situations, so that the network is trained to be robust to variation induced by the unknown aspects.

#### Accuracy with biased sampling

When CKMRnn was both trained and tested on constant-size simulations, the predicted population sizes closely matched true population sizes for all levels of sampling bias (Figure 4a). On average, CKMRnn estimated population sizes within 6.3% of the true population size across all levels of spatial sampling bias. Estimates were unbiased, with mean relative error less that 1.5% in all cases: 0.011, 0.001, and 0.013 for uniform, medium bias, and high bias sampling plans, respectively (Figure 4b, Table 2).

**Figure 4:**
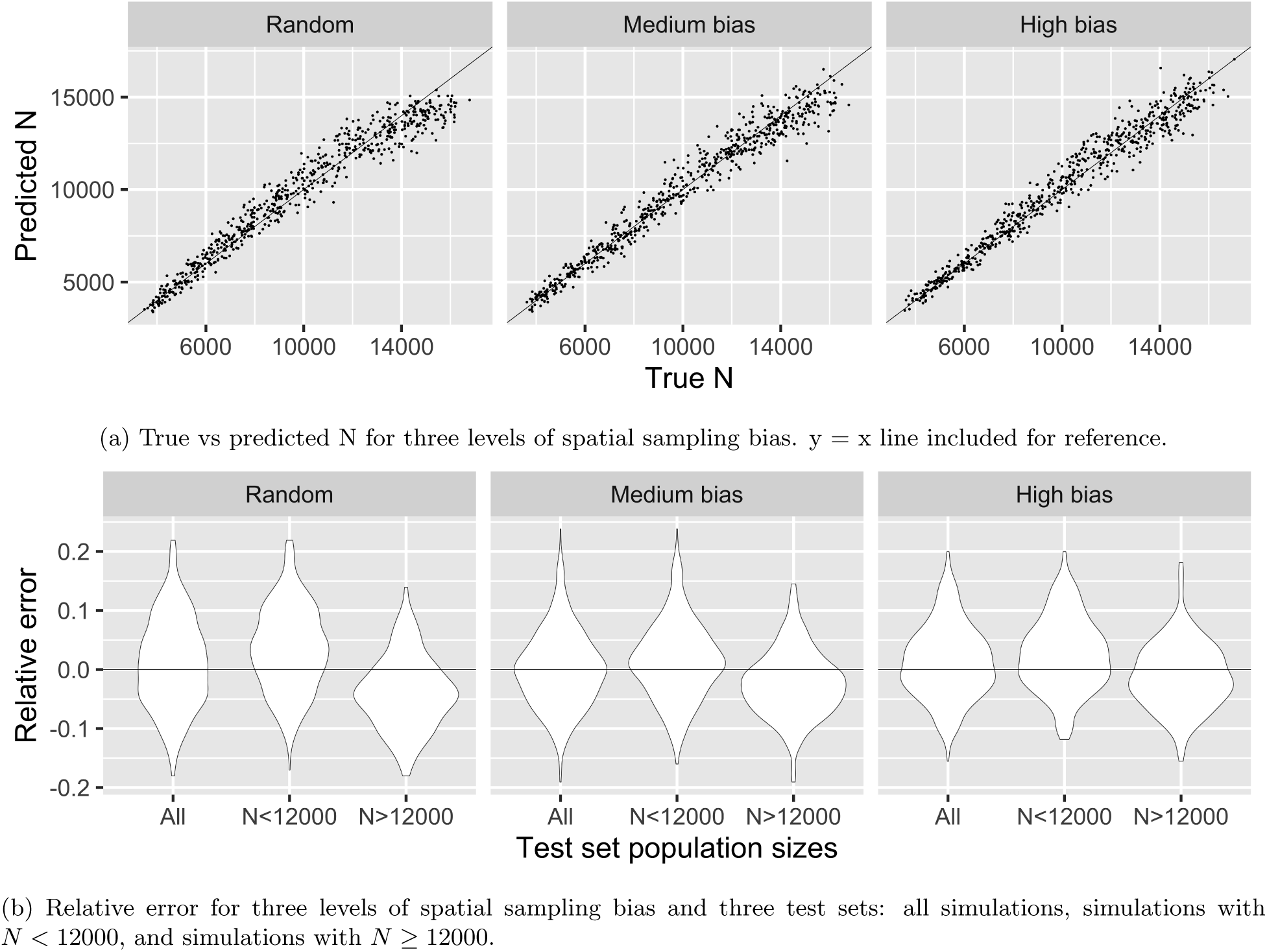
Performance of CKMRnn when trained and tested on simulations with constant population size over time.

**Table 2:** CKMRnn performance with varying levels of spatial sampling bias, as depicted in. **Figure 4a**. Here, the population sizes of both the tested simulations and the simulations used to train the model were constant in time (i.e., no population trend). Relative absolute error is calculated as [true − predicted]*/*true and relative error as (true − predicted) */*true. Separate results are shown for simulations with true *N* above and below 12,000.

| CKMRnn performance with sampling bias |  |  |  |  |  |  |
| --- | --- | --- | --- | --- | --- | --- |
| Spatial sampling bias | Mean relative absolute error (MRAE) | MRAE $N < 12000$ | MRAE $N \geq 12000$ | Mean relative error (MRE) | MRE $N < 12000$ | MRE $N \geq 12000$ |
| uniform | 0.063 | 0.064 | 0.060 | 0.011 | 0.034 | -0.039 |
| medium | 0.051 | 0.052 | 0.049 | 0.001 | 0.014 | -0.024 |
| high | 0.050 | 0.053 | 0.046 | 0.013 | 0.025 | -0.011 |

As true population size increased, estimates appeared to become less accurate and more negatively biased, so we also divided the test set into populations below and above size 12000 and computed relative error for each. Mean relative error (MRE) and mean relative absolute error (MRAE) were about the same for *N <* 12000 and *N* ≥ 12000 but errors were shifted slightly upward for *N <* 12000 and downward for *N* ≥ 12000. This shift was especially apparent for uniform sampling, with MRE for *N <* 12000 of 0.034 and MRE for *N* ≥ 12000 of -0.039. However, errors were still centered near zero. For our simulations we expect estimates to become more negatively biased as population size increases because we are sampling a smaller proportion of the population and will get fewer parent-offspring and half-sibling pairs. Because dispersal is limited, sampling plans with spatial bias tend to sample more related pairs for the same sample size and so this pattern will be less prevalent for biased plans.

#### The effect of misspecified population trend

When CKMRnn was trained on constant size simulations and tested on simulations with increasing or decreasing trend, estimates of current population size were less accurate and more biased (Figure 5). Overall, CKMRnn estimated final population size to within about 20% of the true value for all sampling plans (Table 3). The estimates were positively biased for test simulations with decreasing population size and negatively biased for those with increasing size (Figure 5). This is expected: when population size is decreasing, the number of kin pairs is smaller than expected for a constant size population and so if the trend is not accounted for, estimates will be negatively biased, and vice versa for increasing trend.

**Figure 5:**
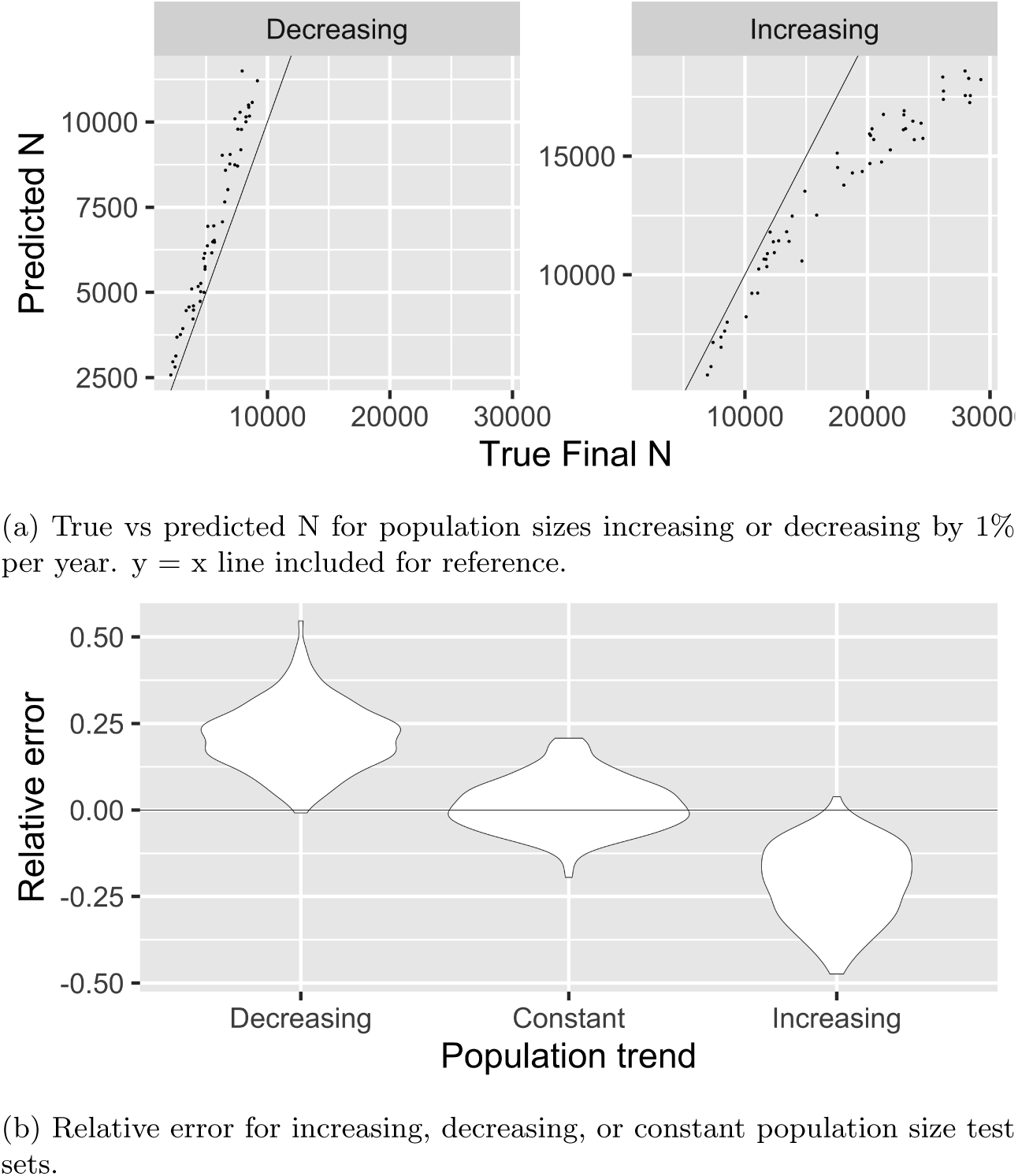
Performance of CKMRnn when trained on simulations with constant population size over time and tested on simulations with increasing, constant, or decreasing population size. All results are for medium spatial sampling bias.

**Table 3:** CKMRnn performance on data with misspecified population trend. The method is trained with simulations of constant population size, but tested on simulations with population size increasing or decreasing by 1% per year.

| **CKMRnn performance for misspecified population trend** |  |  |  |
| --- | --- | --- | --- |
| Population trend | Spatial sampling bias | Mean relative absolute error | Mean relative error |
| decreasing | uniform | 0.206 | 0.206 |
| increasing | uniform | 0.234 | -0.232 |
| decreasing | medium | 0.224 | 0.224 |
| increasing | medium | 0.211 | -0.211 |
| decreasing | high | 0.208 | 0.207 |
| increasing | high | 0.192 | -0.192 |

#### Training for robustness to population trend

Finally, we tested our ability to make CKMRnn robust to unknown population trend by training the network on a set of simulations that included increasing, decreasing, and constant trend populations. We did not provide the network any information about population trend beyond what it could infer from the input images. The range of trends in the training set is conceptually similar to a prior distribution, in that the network is expected to work best when the true trend lies within the “prior” range. We do not ask the network to also infer trend: indeed, the network might be either effectively learning the trend and adjusting accordingly, or learning to use patterns that are unaffected by the trend (or both). The updated network estimated population size to within about 10% of the true value for all population trends and sampling biases (Table 4). This robustness comes at little cost – it was much more accurate than when trend was misspecified but only slightly less accurate than when trend was perfectly specified. Errors still tended to be positive for decreasing trend and negative for increasing trend, but only slightly (Figure 6).

**Figure 6:**
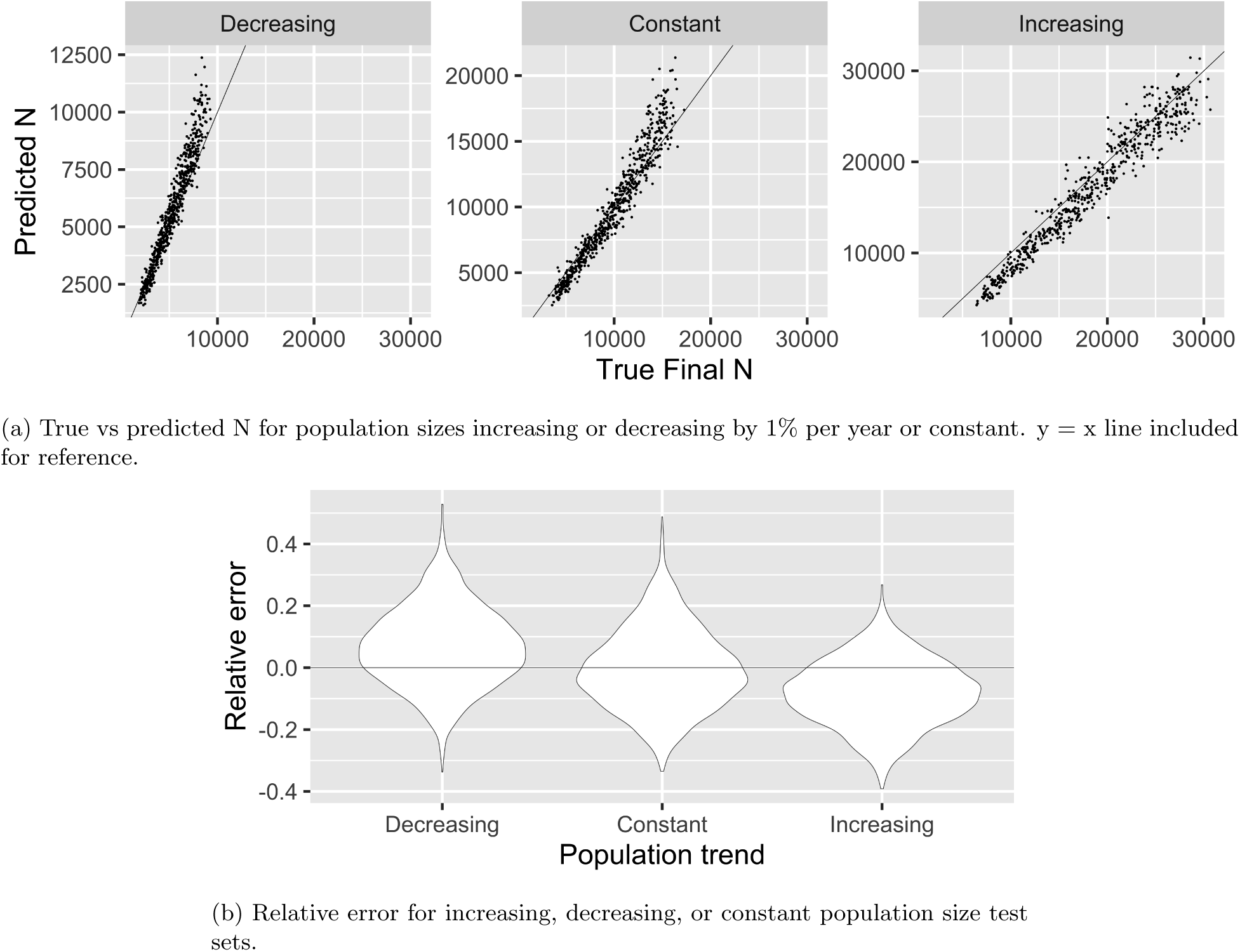
Performance of CKMRnn when trained and tested on simulations with increasing, constant, or decreasing population size. All results are for medium spatial sampling bias.

**Table 4:** CKMRnn performance on datasets with mixed population trends, using a single network trained across a range of trends. Both training and test sets are simulations with population size increasing or decreasing by 1% per year or constant population size.

| **CKMRnn performance for unknown population trend** |  |  |  |
| --- | --- | --- | --- |
| Population trend | Spatial sampling bias | Mean relative absolute error | Mean relative error |
| decreasing | uniform | 0.110 | 0.065 |
| constant | uniform | 0.109 | -0.002 |
| increasing | uniform | 0.116 | -0.093 |
| decreasing | medium | 0.108 | 0.056 |
| constant | medium | 0.099 | -0.020 |
| increasing | medium | 0.109 | -0.085 |
| decreasing | high | 0.122 | 0.078 |
| constant | high | 0.105 | 0.011 |
| increasing | high | 0.099 | -0.064 |

### African elephants

The African elephant dataset contains genetic information from 256 dung piles [17]. These samples are concentrated on the outer edges of Kibale National Park, with few samples from the middle, suggesting that there is strong spatial bias in sampling (Figure 2). The dung samples contain 124 unique individuals, 103 of which were not recaptured, 16 were captured on two different days, 1 on three different days, 3 on four days and 1 on five days. Distances between subsequent recaptures ranged between 0.54 km and 33.94 km, with an average distance of 9.32 km and an average distance per day of 0.79 km. We plotted these recapture histories for input to the neural network by connecting subsequent recaptures of the same individual on different days with line segments, resulting in 47 line segments (Figure 2).

Among the unique individuals, there were 260 parent-offspring pairs. The distance between parent and offspring sampling locations ranged from 0 to 47.12 km, with an average of 17.84 km. Some of the individuals in the pairs were captured multiple times. We plotted these parent-offspring pairs for input to the neural network by treating samples from the same individual on two different days as unique individuals, resulting in 338 line segments, shown in Figure 2.

To validate the method in this system, we first tested the performance of CKMRnn on simulated elephant populations by training the network on 7500 simulations and testing on the remaining 2500. The neural network performed very well on the held out test set. Estimates were unbiased and estimated population size was on average within 12.3% of the true value for population sizes between 100 and 1500 (Figure 7). The variation in estimates increased somewhat as population size increased (Figure 7a). This is expected because sample size relative to population size decreases as population size increases.

**Figure 7:**
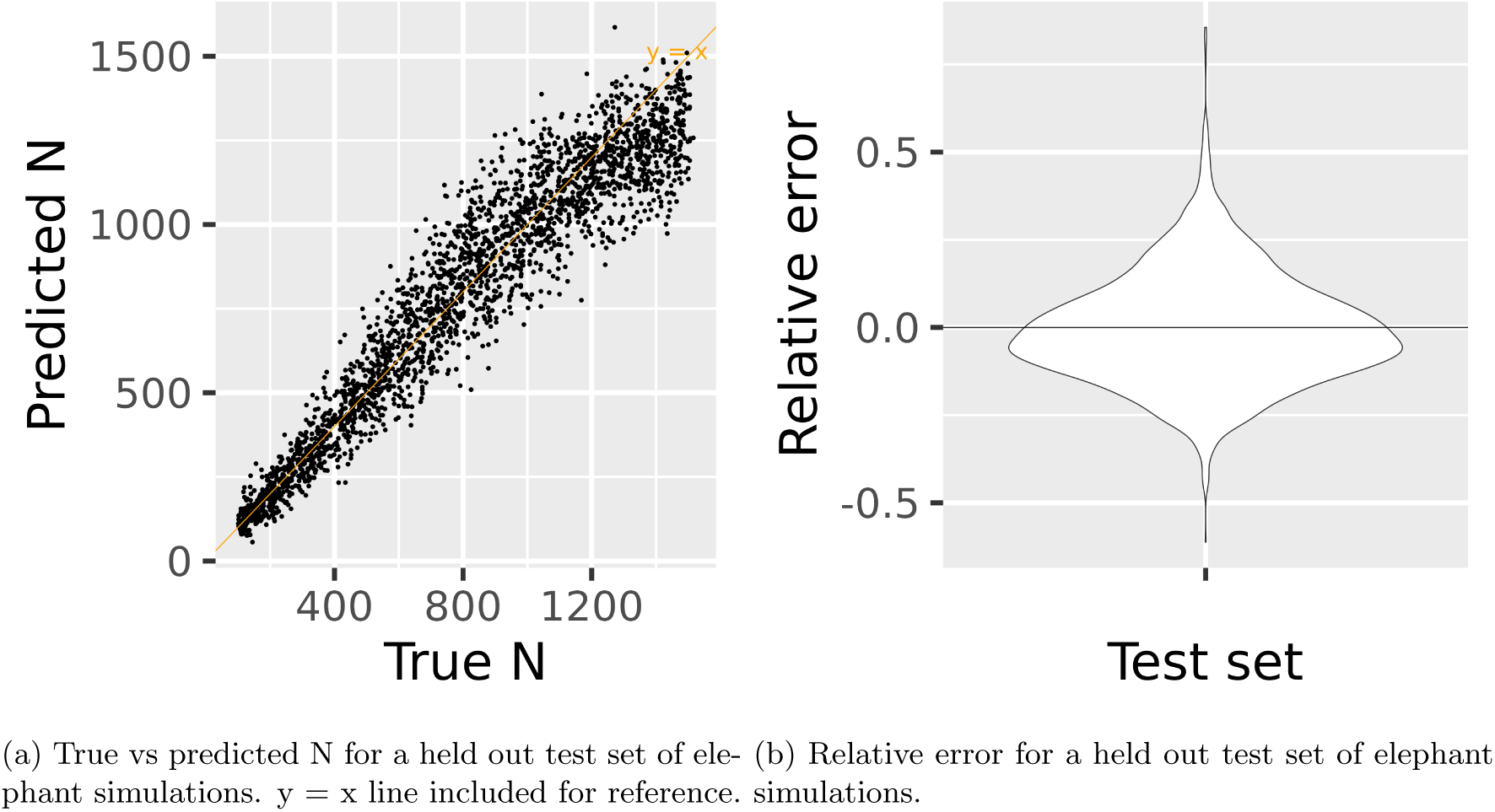
Performance of CKMRnn on elephant simulations.

Finally, we used our trained network to estimate population size in Kibale National Park from images of empirical sampling intensity, recaptures, and parent-offspring pairs, obtaining an estimate of 450 elephants. To produce a confidence interval, we generated 500 parametric bootstrap replicates by simulating 500 populations each with population size 450, then used the network to estimate population size for each replicate to get a bootstrap distribution (Figure 8). The 95% confidence interval based on the distribution is (32, 620).

**Figure 8:**
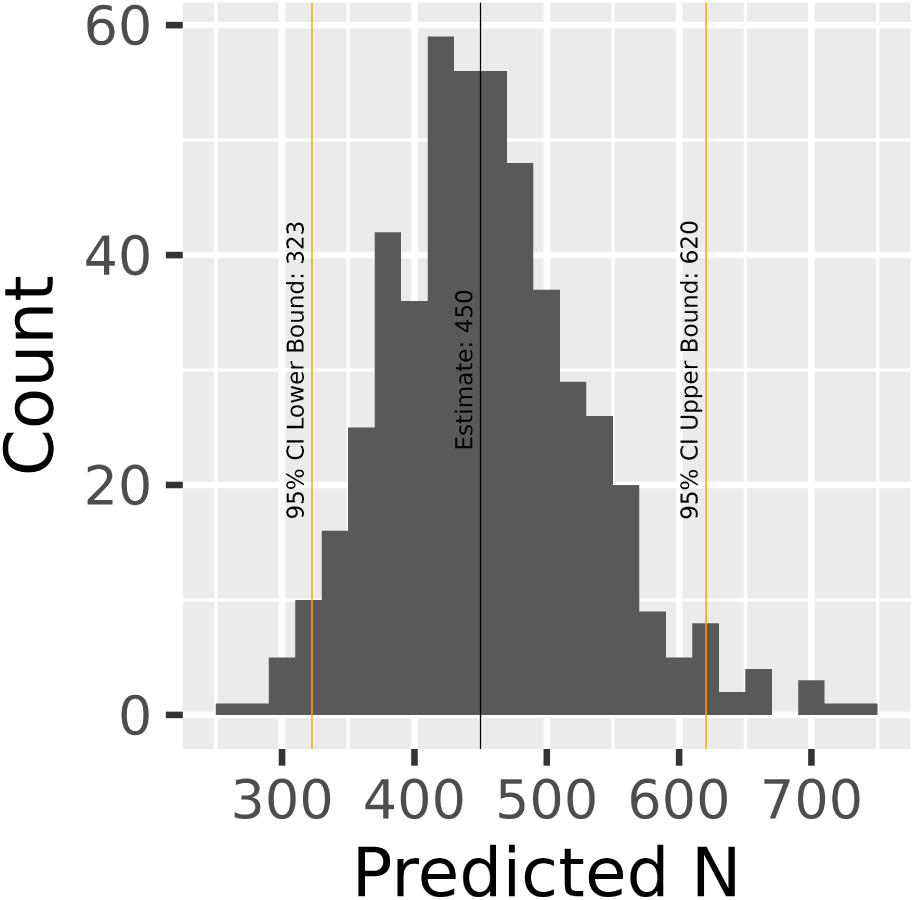
Histogram of parametric bootstrap replicates for population size of African elephants in Kibale National Park. Vertical lines are the point estimate and bounds of the 95% confidence interval.

For comparison, we ran a traditional capture-recapture analysis of the elephant dataset to estimate population size. We used a log-linear model in the R package Rcapture [3]. Based on AIC scores, the best closed population, continuous time model was Chao’s Lower Bound with heterogeneity in capture probability between individuals [32]. The estimated population size was 454.1 with a 95% CI of (316.3, 752.9).

There are two previous estimates of population size in Kibale, a 2021 cut-transect survey estimate of 566 elephants with a 95% CI of 377 to 850 from Daniel, Edward, and Kennedy [11] and the 2025 capture-recapture estimate of 573 elephants with a 95% CI of 410 to 916 from Goodfellow et al. [17].

Our CKMRnn estimate is notably smaller than the estimate of 573 obtained by Goodfellow et al. [17] using the same capture-recapture data, but still falls within the 95% confidence interval of this estimate. This difference in estimates is likely due to differences in the way we accounted for non-independence between sampling locations. Goodfellow et al. [17] accounted for non-independence by filtering the data to include only one capture instance for individuals with multiple captures near the same sampling location. We accounted for non-independence using a much less strict filtering process (only filtering captures of the same individual on the same day), and then accounting for the remaining non-independence through simulation of the sampling plan.

## Discussion

In this paper we develop a spatially explicit close-kin mark-recapture method, CKMRnn, that uses a convolutional neural network (CNN) to estimate population size from maps of sampling intensity, kin pairs, and other relevant information such as recaptures. Our method is accurate and robust, even for populations with limited dispersal and spatially-biased sampling, a situation in which most previous methods are biased [9, 34]. On simulated test populations with spatial population structure and spatially-biased sampling, CKMRnn was unbiased and estimated true population size to within 6% of the true value when population trend was constant over time and to within about 10% when population trend was unknown. CKMRnn also performs well on empirical data. We used CKMRnn to estimate population size of elephants in Kibale National Park in Uganda from recaptures and parent-offspring pairs. We estimated that there are around 450 elephants in the park, with a 95% confidence interval of 323 to 620. The estimate very closely agrees with estimated population size using a traditional capture-recapture method and has a 32% narrower confidence interval.

CKMRnn has several advantages over previously-developed spatial and simulation-based CKMR methods. For instance, the spatial methods developed by Sévêque et al. [34] use a pseudo-likelihood-based approach to account for spatially-biased sampling. This approach is limited to simple models of spatial population dynamics. For example, it assumes that dispersal distances are independent between individuals and that the sampling location of a female is the initial location of the offspring. In addition, the pseudo-likelihood approach assumes sampled pairs are independent and so is best suited to systems where population size is large relative to sample size. Relaxing such assumptions is difficult in the likelihood setting because analytical expressions for observing kin pairs quickly become hard or impossible to write down. Because CKMRnn instead relies upon simulation, the complexity of the model is limited only by our ability to simulate it. Our elephant example demonstrates how this flexibility allows us to apply CKMR to populations that violate the assumptions made by likelihood methods. In the small elephant population, sampled pairs are not independent, and we are able to account for this non-independence by simulating it. Juvenile elephants stay with their mothers for many years before dispersing, and we are able to account for this pattern by explicitly simulating the movements of mothers and juveniles. In the future, we plan to extend CKMRnn to account for even more complex dynamics such as correlated movement of closely related individuals in kin groups.

Other researchers have also implemented simulation-based methods. For instance, the simulation-based methods developed by Conn [8] use Approximate Bayesian Computation (ABC) to infer population size. The observed data for this ABC method is counts of kin pairs grouped by relevant information such as age class. This ABC method works well, but it does not directly account for spatial information. Each sampled individual has a unique location on the landscape, and so when pairs of individuals are aggregated into groups to use as observed data for the ABC method, these unique locations are lost. Our CNN, which relies on an image-based input format that includes the spatial locations of all samples, thus has the potential to take advantage of spatial information.

Currently, there are three major limitations of CKMRnn: (1) it does not explicitly include information about time; (2) it estimates a single population density rather than a map of density that varies across the landscape, and (3) it assumes that kin identification is without error. We believe that CKMRnn could be extended to relax these assumptions, but will leave this for future work.

In pseudo-likelihood methods, time is included in the model through ages of sampled individuals and through information about sampling time. In these methods, both parent-offspring and half-sibling pair probabilities are computed based on the birth year of the potential offspring or the birth years of the potential half-siblings [6], and so ages of sampled individuals are required and error in aging can cause bias in abundance estimates [31, 38]. Observing ages and sampling times also allows these methods to estimate parameters such as survival and population trend. CKMRnn does not use ages of individuals as input to the network, and assumes that the age structure of the population is known (since this is used in setting up the simulation).

In Kibale, the available samples are from only one year and age of the sampled individuals is unknown, and so we do not have much information about parameters that depend on time, such as survival or population trend. Elephants are also fairly well studied, and so we have prior information on the age structure and survival dynamics of elephants. However, for populations that are sampled yearly such as through hunter harvests, or for populations that are not as well-studied as elephants and so age structure and survival is unknown, we would want to incorporate age and time information. Future work to include time might vary age-dependent survival parameters in the training simulations and/or adapt the network architecture to take as input age and sample time information. Such networks might also estimate parameters such as survival and trend in addition to population size.

Our applications here have assumed that the population is closed and that population density is roughly constant across the landscape. Most populations, including the Kibale elephants, do not meet these assumptions. Despite these assumptions, CKMRnn was able to get reasonably accurate estimates for Kibale, however, for populations with greater variation in density across the landscape or higher levels of immigration and emigration, we would likely need to relax these assumptions. For example, in populations without clear boundaries, estimating a single, closed population size is not meaningful [33]. In addition, population density that varies across the landscape influences patterns of kin pairs and needs to be accounted for. Previous methods such as that from Sévêque et al. [34] account for varying density by assuming that the relative density of individuals across the landscape is known from auxiliary data such as camera traps. If auxiliary data was available, this approach could easily be incorporated into CKMRnn by either holding it fixed (as for our simulated elephants, who only live within the park), or adding a map of relative density to the training simulations. If auxiliary data were not available, one could use our approach by varying maps of relative density in the training simulations, which we would expect simply to increase uncertainty in the estimated population sizes. One especially exciting future direction is to extend CKMRnn to directly estimate ecological parameters. For example, we can add a relationship between forest cover and population density to our simulation, and then estimate the parameters of this relationship. Spatially explicit capture-recapture models are already able to incorporate these ecological parameters [33].

Finally, in this paper we assumed that there was no uncertainty in identifying kin pairs. Bravington, Skaug, and Anderson [6] describe two ways of accounting for uncertainty, first, by adding a false negative rate for kin pair estimation to the model and then estimating this rate along with other parameters, and second, by treating kinship as a latent variable and using genetic data as the observation. Both of these approaches could be adapted to account for uncertainty in identifying kin pairs in CKMRnn. For example, we can add a false negative rate for each kin relationship to the training simulations. Or, for populations with significant inbreeding or population structure that influences accuracy of kin identification, we can simulate genotypes of individuals, and then use these genotypes to estimate kin and generate training images. It might even be possible to follow previous methods that use neural networks to estimate other population parameters [35, 36, 37] and use genotypes directly as input to the network.

A natural question in application of CKMRnn is: how accurate does the simulation need to be? Implementing an individual-based spatial simulation can be daunting, as it requires specification of many behaviors, such as mate-finding and dispersal, that may not be well-understood. For a recent, comprehensive guide to implementing spatial simulations, see Chevy et al. [7]. Clearly, no simulation can perfectly capture the precise demographic dynamics of a given species, so some degree of approximation is necessary. This is not unique to simulation-based inference: similarly, any analytical method makes a variety of assumptions about the underlying model (and, these assumptions are often less obvious). As we’ve demonstrated here using population trend, robustness to uncertainty in model parameters can be trained directly into CKMRnn by simulating across a range of parameters in training data. Which aspects of demography are important to model and how model uncertainty affects accuracy of estimates are empirical questions that will depend on the system being studied.

Some aspects of uncertainty are, however, unavoidable. Similar to how individual heterogeneity in capture probability leads to nonidentifiability in mark-recapture estimates [32], variation in offspring number distribution can lead to nonidentifiability in CKMR estimates of population size, at least for nonspatial models [22, 25]. The distinction is related to “effective” versus “census” population size, *N_e_*versus *N* [41]. One definition of *N_e_*is in terms of the proportion of pairs that are siblings, and so two populations having the same sibling density may have the same *N_e_* but different *N*. To the degree that a CKMR method relies on sibling density, therefore, it may be *N_e_* rather than *N* that is estimable and so different models with the same *N_e_*but different *N* may be difficult or impossible to distinguish. CKMRnn predicts *N*, not *N_e_*(because that is what it is given for training data), but in some situations *N_e_*might be more accurately estimable. Information about *N* beyond *N_e_*comes from two sources: from “prior” information about the demographic model (i.e., how the simulation is coded), and from time-varying data (e.g., mark-recaptures or parent-offspring relationships). Better understanding of these aspects could improve future study design and interpretation.

Many parts of the CKMRnn pipeline are specific to the population being studied, and so we do not provide an R or python package that can be directly applied to a new system. Much of the code we used to run CKMRnn on the Kibale elephants can be adapted for a new system, and we provide this code as well as instructions for running it in the CKMRnn GitHub repository. The most challenging step in applying CKMRnn to a new system will likely be writing a reasonably accurate simulation of the population and sampling plan. The pipelines we provide for finding and visualizing kin pairs and recaptures and generating training simulations could be directly used with very few changes. We also provide a pipeline for training the network and obtaining estimates and parametric bootstrap confidence intervals. This pipeline can also be directly used with few changes, but requires access to a GPU.

In summary, we have found that our new method, CKMRnn, has the potential to improve monitoring for many terrestrial species, which often have population structure and spatially-biased sampling. We expect CKMRnn to be especially useful for populations such as the Kibale elephants where individuals are elusive and hard to capture.

## Acknowledgements

We thank the Kern-Ralph co-lab for input and comments on the project. Funding was provided by NIH awards R35148253, R01HG010774, and R01HG012473.

## Data Availability

Code and data are available at https://github.com/giliapatterson/CKMRnn.

## Notes

https://github.com/giliapatterson/CKMRnn

### Competing Interest Statement

The authors have declared no competing interest.

